# Description of three new species of predatory Genus *Hexacentrus* (Orthoptera: Tettigoniidae) from India, with bioacoustic and morphological characterization

**DOI:** 10.1101/2023.01.24.525294

**Authors:** Aarini Ghosh, Ranjana Jaiswara, Monaal, Shagun Sabharwal, Vivek Dasoju, Anubhab Bhattacharjee, Bittu Kaveri Rajaraman

## Abstract

*Hexacentrus* is a genus of predatory katydids. In India, the genus *Hexacentrus* is represented by 7 species of which 6 are morphologically characterized while one is only acoustically characterized. In this study, we describe three new species under the genus from India based on detailed morphological, stridulatory, and acoustic features.

## Introduction

*Hexacentrus* species (Hexacentrinae, Tettigoniidae, Orthoptera) are small to moderate-sized predatory katydids (Heller 1986), found largely in Africa, Asia, Australia, and the Pacific (Farooqi & Usmani 2018; Grzywacz *et al*. 2021). *Hexacentrus* species are largely green with brown segments and are well camouflaged in bushes. The male tegmen is modified for sound production while the female tegmen is devoid of these specializations. *Hexacentrus* can be distinguished by their unique leaf-shaped wings, and six pairs of spurs in the fore and mid tibia.

*Hexacentrus* is one of 13 Hexacentrine genera, and there are 28 known species of *Hexacentrus*, one of which has two subspecies (Cigliano *et al*. 2017; Chen *et al*. 2021). Some additional morphologically cryptic species from the genus *Hexacentrus* have been identified using DNA barcoding (Chen *et al*. 2021; Guo *et al*. 2016; Kim *et al*. 2020). India is home to 6 morphologically characterized species of *Hexacentrus* and a seventh for which only an acoustic description is available (Table 1). Of these, the first description of *Hexacentrus* was of *H. unicolor* Serville, 1831, and this along with *H. mundus* Walker, 1869 has been recorded multiple times from the north-eastern part of India by the Zoological Survey of India (ZSI) and in the Orthoptera species file (Shishodia 2006; Shishodia *et al*. 2010; Ingrisch & Shishodia, 1998; Srinivasan & Prabakar 2012; Cigliano *et al*. 2017). *H. bifurcata* was discovered by Farooqi & Usmani (2018) alongside a new record of sympatric *H. japonicus* Karny, 1907, and sympatric *H. expansus* (Wang and Shi, 2005; Farooqui & Usmani 2019) from Uttar Pradesh in the northern part of India. *H. major* Redetenbacher 1891 was recently found and acoustically described from the Bhagwan Mahavir Wildlife sanctuary of Goa (Tiwari *et al*. 2022). Tiwari *et al*. (2022) also documented an acoustic signal from a morphologically partly characterized species of *Hexacentrus* from the Hollongapar Gibbon Wildlife sanctuary of Assam. Other than this uncharacterized caller which they call *H. sp, H*.*major* and *H*.*bifurcata*, which are found only in India, the remaining species are widely distributed in South East Asia (Chen *et al*. 2021; Guo *et al*. 2016; Ichikawa *et al*. 2006; Gorochov & Sliwa 1999; Panhwar *et al*. 2014; Storozhenko *et al*. 2015). Acoustic data are available only for three of these other *Hexacentrus* species: *H. mundus, H. major*, and *H. unicolor*. In our study, we found three more species of *Hexacentrus* in India that differ morphologically and acoustically from these previously described species.

**Table 1.** The list of *Hexacentrus* species found in India (Shishodia (2006); Shishodia *et al*. (2010); Ingrisch & Shishodia (2000); Srinivasan & Prabakar (2012), Tiwari *et al*. 2022)

| Species | Distribution recorded in India | Distribution recorded in other countries |
| --- | --- | --- |
| *Hexacentrus unicolor* Serville, 1831 | Nagaland, Mizoram, Andaman & Nicobar, Manipur, Himachal Pradesh, Assam | South-East Asian countries |
| *Hexacentrus major* Redtenbacher, 1891 | Orissa, Goa | Not recorded. |
| *Hexacentrus bifurcatus* Farooqi & Usmani, 2018 | Uttar Pradesh | Not recorded. |
| *Hexacentrus japonicus* Karny, 1907 | Uttar Pradesh | South-East Asian countries |
| *Hexacentrus expansus* Wang & Shi, 2005 | Uttar Pradesh | China |
| *Hexacentrus mundus* Walker, 1869 | Rajasthan, Assam, Meghalaya | Borneo, Japan, Philippines, Moluccas |
| *Hexacentrus* sp. (Tiwari & Diwakar, 2022) | Assam | Not recorded |

**Table 2.** Measurements of male and female ***H. khasiensis*** Ghosh, Jaiswara & Rajaraman **sp. nov**. Abbreviations listed in material and method section. Values in mm.

| Males | FW_L | BL_FWL | PH | PL | FI_L | TI_L | FIII_L | TIII_L |  |
| --- | --- | --- | --- | --- | --- | --- | --- | --- | --- |
| Holotype | 21.4 | 30 | 2.9 | 6.5 | 6.5 | 7.3 | 17.4 | 17.3 |  |
| M_1<br>paratype<br>dry | 21.3 | 30 | 4 | 6.4 | 6.7 | 5.7 | 15.8 | 15.2 |  |
| M_2<br>paratype<br>dry | 21.5 | 32 | 3.9 | 6.5 | 6.5 | 7.5 | 16 | 17 |  |
| M_3<br>paratype<br>wet | 21.3 | 31 | 3.1 | 6.3 | 6.3 | 8 | 15.7 | 17.3 |  |
| M_4<br>paratype<br>wet | 23.4 | 35 | 3.7 | 7 | 6.3 | 7.8 | 17.8 | 17.8 |  |
| Females | BL | FW_L | Ovi_L | PH | PL | FI_L | TI_L | FIII_L | TIII_L |
| Allotype | 25.3 | 19.4 | 14 | 3.5 | 5.4 | 6.9 | 7.6 | 17.3 | 18.5 |
| **F_1**<br>**paratype**<br>**dry** | 23.5 | 20 | 11.32 | 4.4 | 5.2 | 6.8 | 8.7 | 18 | 18.3 |
| **F_2**<br>**paratype**<br>**wet** | 26.5 | 20.2 | 14.3 | 3.2 | 5.3 | 6.4 | 7.1 | 17 | 17.4 |
| **F_3**<br>**paratype**<br>**wet** | 22.8 | 19.5 | 13.6 | 3.9 | 5.5 | 4.2 | 4.8 | 9.9 | 11 |

**Table 3.** Measurements of male and female ***H. ashoka*** Ghosh, Jaiswara & Rajaraman, **sp. nov**. Abbreviations listed in material and method section. Values in mm.

| **Males** | **BL** | **FW_L** | **BL_FWL** | **PH** | **PL** | **FI_L** | **TI_L** | **FIII_L** | **TIII_L** |
| --- | --- | --- | --- | --- | --- | --- | --- | --- | --- |
| **Holotype** | 23.3 | 33 | 42 | 3.6 | 6.9 | 6.6 | 8.7 | 16.7 | 20 |
| **M_1**<br>**Paratype**<br>**dry** | 23.3 | 32 | 42 | 3.7 | 7.5 | 6 | 8.2 | 16.5 | 19.5 |
| **M_2**<br>**Paratype**<br>**dry** | 22.3 | 33 | 41 | 4.3 | 7.5 | 7.3 | 8.1 | 17 | 19.4 |
| **Females** | **BL** | **FW_L** | **Ovi_L** | **PH** | **PL** | **FI_L** | **TI_L** | **FIII_L** | **TIII_L** |
| **Allotype** | 25 | 30 | 14 | 4.5 | 5.6 | 5.8 | 7.6 | 19.4 | 16 |
| **F_1**<br>**paratype**<br>**dry** | 22.63 | 30 | 14.4 | 4 | 6 | 6.8 | 10.1 | 20.2 | 20.5 |
| **F_2**<br>**paratype**<br>**wet** | 21.2 | 30 | 14.7 | 4.7 | 5.4 | 7.2 | 8.5 | 20.5 | 20.6 |
| **F_3**<br>**paratype**<br>**dry** | 23 | 30 | 14.1 | 3.8 | 5.3 | 6.9 | 7.7 | 19 | 19.1 |

**Table 4.** Measurements of male and female ***H. tiddae*** Ghosh, Jaiswara, Monaal & Rajaraman, **sp. nov**. Abbreviations listed in material and method section. Values in mm.

| **Males** | **BL** | **FW_L** | **BL_FWL** | **PH** | **PL** | **FI_L** | **TI_L** | **FIII_L** | **TIII_L** |
| --- | --- | --- | --- | --- | --- | --- | --- | --- | --- |
| **Holotype** | 27 | 36 | 45 | 3.4 | 6.5 | 6.4 | 7.6 | 16.7 | 16.5 |
| **M_1**<br>**paratype**<br>**dry** | 22.7 | 34 | 42 | 3.3 | 6.5 | 6.5 | 6.7 | 15.1 | 16.2 |

## Methods

### Study Sites

Our first study site was in the forest area around and inside the North-eastern Hill University (NEHU) campus of Shillong, Meghalaya (Fig. 1), India at an altitude of 1390-1410m a.s.l., GPS location-25° 36’ 59.76” N; 91° 54’ 2.52” E. This forest area mostly consists of sub-tropical bushes and coniferous trees. In the post-monsoon season (August-September) the precipitation rate was 500-700 mm, and the average temperature and humidity were 20°C and 94% respectively.

**Figure 1.**
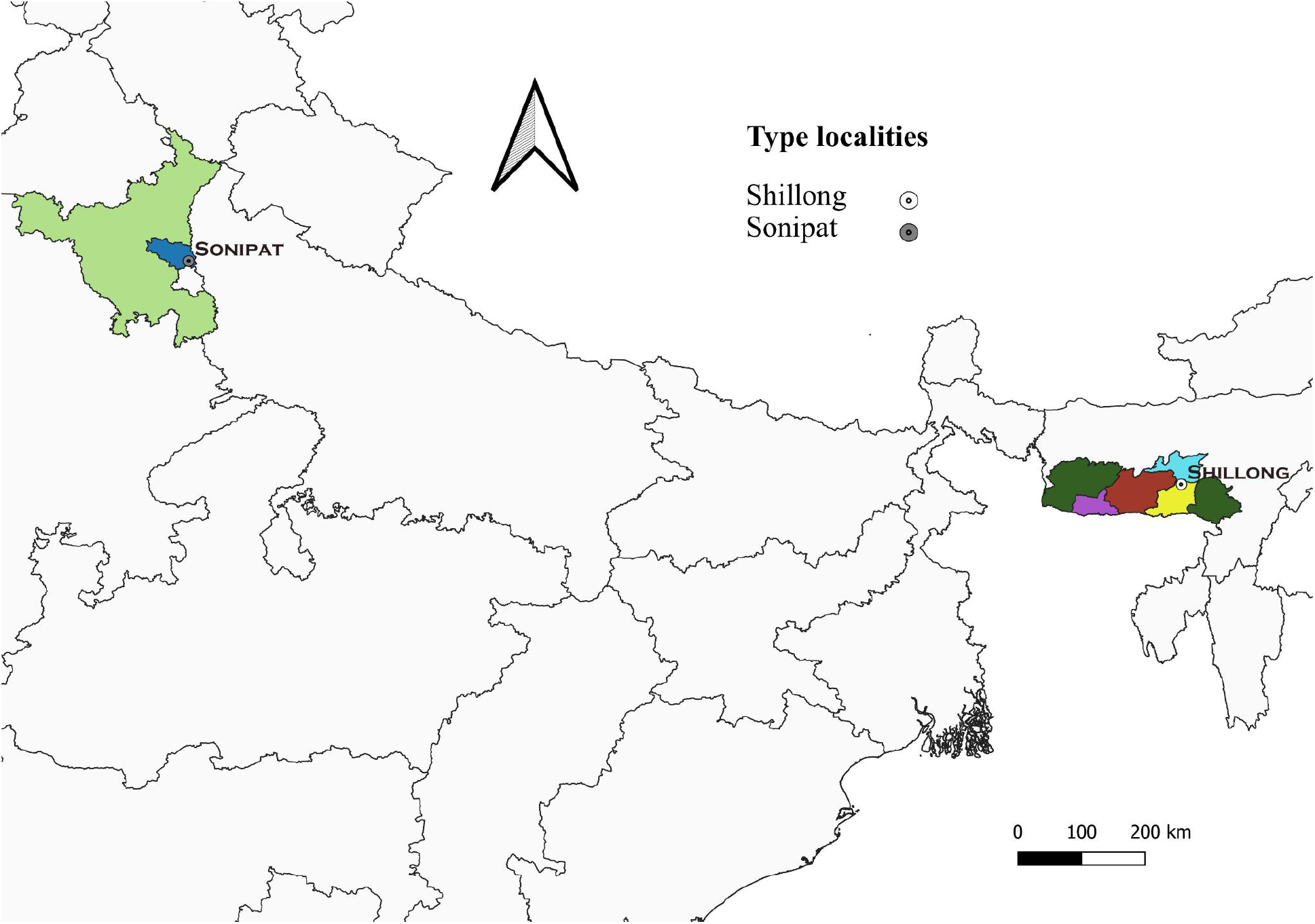
India map showing distributions for all three species including their type localities. The light green colour shows Haryana state, and the dark green colour shows Meghalaya state with other colours highlighting the distribution of ***H. ashoka*** and ***H. tiddae*** in Sonipat district (dark blue); ***H. khasiensis*** distribution in the East-Khasi hills (yellow), West-Khasi hills (red), Ri-Bhoi (sky blue), South Garo hills (purple).

**Figure 2.**
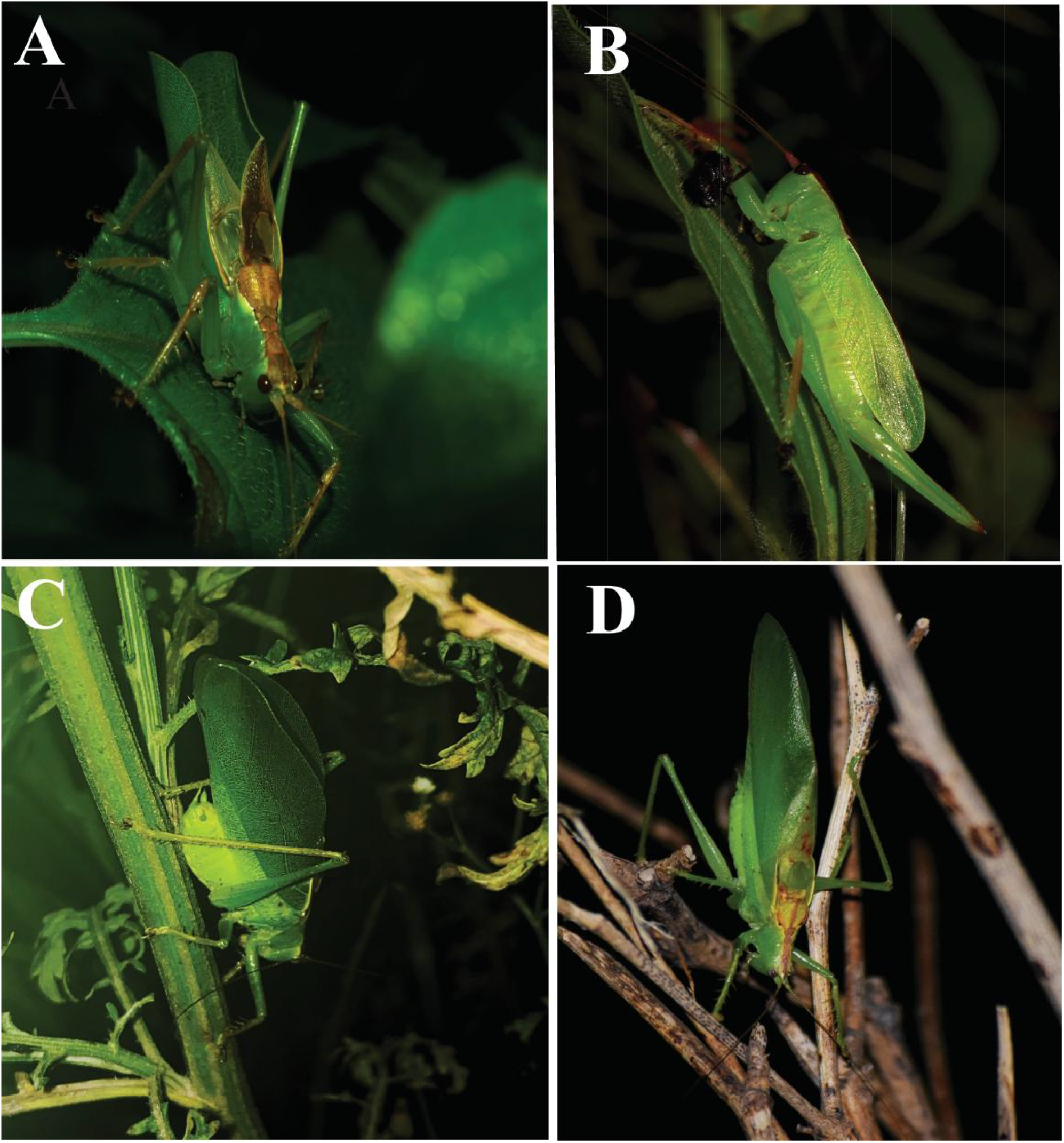
Picture of all 3 species in Wild- ***H. khasiensis*** male (A) and female (B), ***H. ashoka*** male (C), ***H. tiddae*** male (D).

Our second study site was in the bushy area around the Ashoka University Campus, Sonipat, Haryana (Fig. 1), India at an altitude of 315m a.s.l., GPS location-28° 56’ 48.78” N;77° 6’ 5.19” E. This area was mostly covered with thorny bushes and small patches of dry deciduous forest. During the post-monsoon season (August-September) the precipitation rate was 150-200 mm and the average temperature and humidity 30°C and 70% respectively.

### Specimen collection and preservation

*Hexacentrus* are largely found on bushes, from lower to middle vertical strata. They inhabit a variety of habitats, from deciduous forests to thorny scrub areas, to dense sub-tropical forests. Male specimens were collected by focally locating the call positions in the bushes. Females were collected by visual search of the bushes near the calling male. Collections were done from mid-July to September of 2019 and 2021 between 7:00 pm-12:00 midnight. All specimens for taxonomic identification were dry preserved. A few male and female specimens from each species were preserved in 70% ethyl alcohol for morphometric measurements at the Neuroethology lab of Ashoka University.

### Acoustic recording and analyses

Male calling songs of all three species were recorded in the wild using a Marantz PMD661 MK II recorder at 44.1 kHz sampling rate and Pettersson M500 USB ultrasound microphone mounted on a laptop (NOKIA) at 500 kHz sampling rate. The recording was done from 5 feet (60 inches) distance from the caller to avoid clipping. The ambient temperature and humidity of each day were recorded by a digital hygrometer (STORE99 WHDZ HTC-1). Live adult males were carried back to the lab whenever possible to record their call inside a soundproof arena in the Neuroethology lab, at Ashoka University, Sonipat, Haryana, India.

All the temporal and structural parameters of acoustic recordings were analysed using Bat sound 3.31 and Audacity software Version 3.0.5. Power spectrum analysis and spectrograms were made in Bat Sound standard sound analysis software 3.31.

### Morphological structures and measurements

Morphological structures were studied under a stereozoom microscope, Leica S9i (Leica Microsystems, Germany). Measurements of type series of all three species were done using the analysis tool of Leica Application Suite version LAS V4.13.0 installed on a computer running on Windows 10 Operating system in the Department of Zoology, Panjab University, Chandigarh, India and in the Department of Biology, Ashoka University, Haryana.

### Imaging

Scanning Electron microscope imaging of stridulatory teeth was done at 1–2 kV resolution using JSM-6100 (JEOL, Japan) in the Department of Sophisticated Analytical Instrumentation Facility, CIL and UCIM, Panjab University, Chandigarh, India.

### Terminologies

Terminologies of acoustic parameters used are described here. The peak frequency (PF) of the call is the frequency with the highest energy amplitude. We use the term fundamental frequency, defined as the first peak in the spectrum, in case of calls that show harmonics of the fundamental frequency. Bandwidth frequency (BW) is the range of frequency spectrum measured -20dB from the amplitude at peak frequency. Syllables are the smallest unit of the call. Syllable Duration (SD) is the duration of each syllable and the time between two syllables is called the Inter-Syllable interval (ISI). The syllable repetition rate (SRR) is the inverse of the Syllable period. A trill is a series of continuous syllables with no higher superstructure, while a chirp is a series of syllables that lasts for a period called the Chirp Duration (CD) that repeats with a defined Inter-Chirp interval (ICI). The number of syllables per chirp is denoted as (S/C). The term Echeme has been proposed as the highest unit of call structure, useful for more complex call types which may have combinations of chirps and trills.

### Abbreviations

#### General morphology

FI, FII, FIII, fore, median, hind femur; FW, forewing; TI, TII, TIII, fore, median, hind tibia; DD, dorsal disc of pronotum; LL, lateral lobe of pronotum; BL, body length; PH, PL, pronotum height and pronotum length; Ovi_L, ovipositor length; FW_L, forewing length; FI_L, fore femur length; TI_L, fore tibia length; FIII_L, hind femur length; TIII_L, hind tibia length.

#### Tegminal venation

CuA, anterior cubitus; CuA1, CuA2: first, second branches of CuA; CuP, posterior cubitus; M, median vein.

#### Depositories

ZSI: Zoological Survey of India, Kolkata, India.

BNHS: Bombay Natural History Society, Mumbai, India.

AU: Ashoka University, Sonipat, Haryana, India

## Results

### Systematics

Order-Orthoptera

Superfamily-Tettigonioidea Krauss, 1902

Family-Tettigoniidae Krauss, 1902

Subfamily-Hexacentrinae Karny, 1925

### Genus *Hexacentrus* Serville, 1831

#### Type species

*Hexacentrus unicolor* Serville, 1831, by original monotypy.

#### Distribution

Asia, Africa, and Australia.

#### Diagnosis

Same as Serville, 1831; Jian-Feng & Fu-Ming, 2005

### *Hexacentrus khasiensis* Ghosh, Jaiswara & Rajaraman sp. nov

#### Type material

*Holotype*, **♂, INDIA:** Meghalaya, East Khasi Hills, Shillong, ∼1400 m a.s.l., 01.IX.2021, 25° 36’ 59.76” N; 91° 54’ 2.52” E, AG_SH1_M6 (ZSI). *Allotype*, **♀**, AG_SH1_F1, same locality as holotype (ZSI). *Paratypes* (4**♂**, 3**♀**), same information as holotype, VIII-IX.2021: 2**♂ (**AG_SH1_M1, AG_SH1_M2) 2**♂** ex-AG_SH1_M3 (BNHS), ex-AG_SH1_M4 (BNHS); 2**♀ (**AG_SH1_F2, AG_SH1_F3), 1**♀** ex-AG_SH1_F4 (BNHS).

#### Type locality

East Khasi Hill, Shillong, Meghalaya, India (Fig. 1).

#### Distribution

In addition to type locality, this new species has been heard by the research team in different parts of the Khasi hills including the Ri-Bhoi and West Khasi Hill districts of Meghalaya. The same call type has been recorded by the researcher in Balpakram National Park in the South Garo hill district of Meghalaya (Fig. 1).

#### Etymology

The species is named after the Khasi hill of Meghalaya. The species is first time discovered and heard in the parts of the Khasi hills.

#### Diagnosis

The species is somewhat similar to *H. formosanus* Chen & He, 2021 with respect to pronotum markings on the dorsal disc however it differs in several aspects. In *H. formosanus* the lateral margin of the pronotum is straight but in *H. khasiensis* **sp. nov**. it is slightly concave. Teeth in the middle region of the stridulatory file in *H. khasiensis* **sp. nov**. are much more wide than the lateral teeth. Stridulatory teeth are more in number in *H. khasiensis* **sp. nov**. Apical third of anal cerci in *H. khasiensis* **sp. nov**. is longer and sub genital plate styli more slender. Female ovipositor is much more slender.

### Description

**Male**; Body medium sized and slender (Fig. 3A-B). **Head**. Fastigium of vertex triangular, narrow, laterally compressed with median longitudinal sulcus. Apex is narrow, tapering forward, and rounded in lateral view (Fig. 3C). The base of fastigium is 1.25 times as wide as the scapus. The fastigium of vertex is separated from the fastigium of the fons by a furrow. Eyes globular. **Pronotum**. The pronotum is saddle-shaped, longer than broad, and expanded posteriorly. The anterior margin is straight. The posterior margin is slightly concave and the pronotal disc is flat expanded and rounded in the posterior region (Fig. 3C). An hourglass-like band covers the whole pronotum. The median carina is faintly visible. Slightly depressed behind the first transverse sulcus; with three transverse sulci. A ‘V-shaped sulcus is present between the first and second sulci (Fig. 3C). The second and third transverse sulci are restricted to disk only. The ventral margin of the lateral lobe inclines downwards and is longer than high, humeral sinus absent. The anterior angle is obtuse, and the posterior angle is round in shape from the lateral side. The posterior part is a straight slope, and the anterior part is straight downward (Fig. 3E). Thoracic auditory spiracle large and oval, slightly hidden under the lateral lobe of the pronotum. **Legs**. Fore coxae with a forward outward projecting spine. FI dorsally unarmed and ventrally armed with 3-4 sub apical spurs present on inner margin with 4-5 spines present in between and outer margin bearing 4-9 spines, no sub apical spurs. TI dorsally unarmed and ventrally armed with 6 long subapical spurs inwardly bent, first sub apical spur slightly shorter than 2^nd^ sub apical spur and then decreasing in length apically on both sides. TII dorsally armed with two subapical spurs, ventrally with 6 subapical spurs on inner and outer margins. TIII dorsally armed with 22-34 spines on the inner margin and 20-33 spines on outer margin, ventrally armed with 11-15 and 10-13 subapical spurs variable in length and density on outer and inner margin respectively.

**Figure 3.**
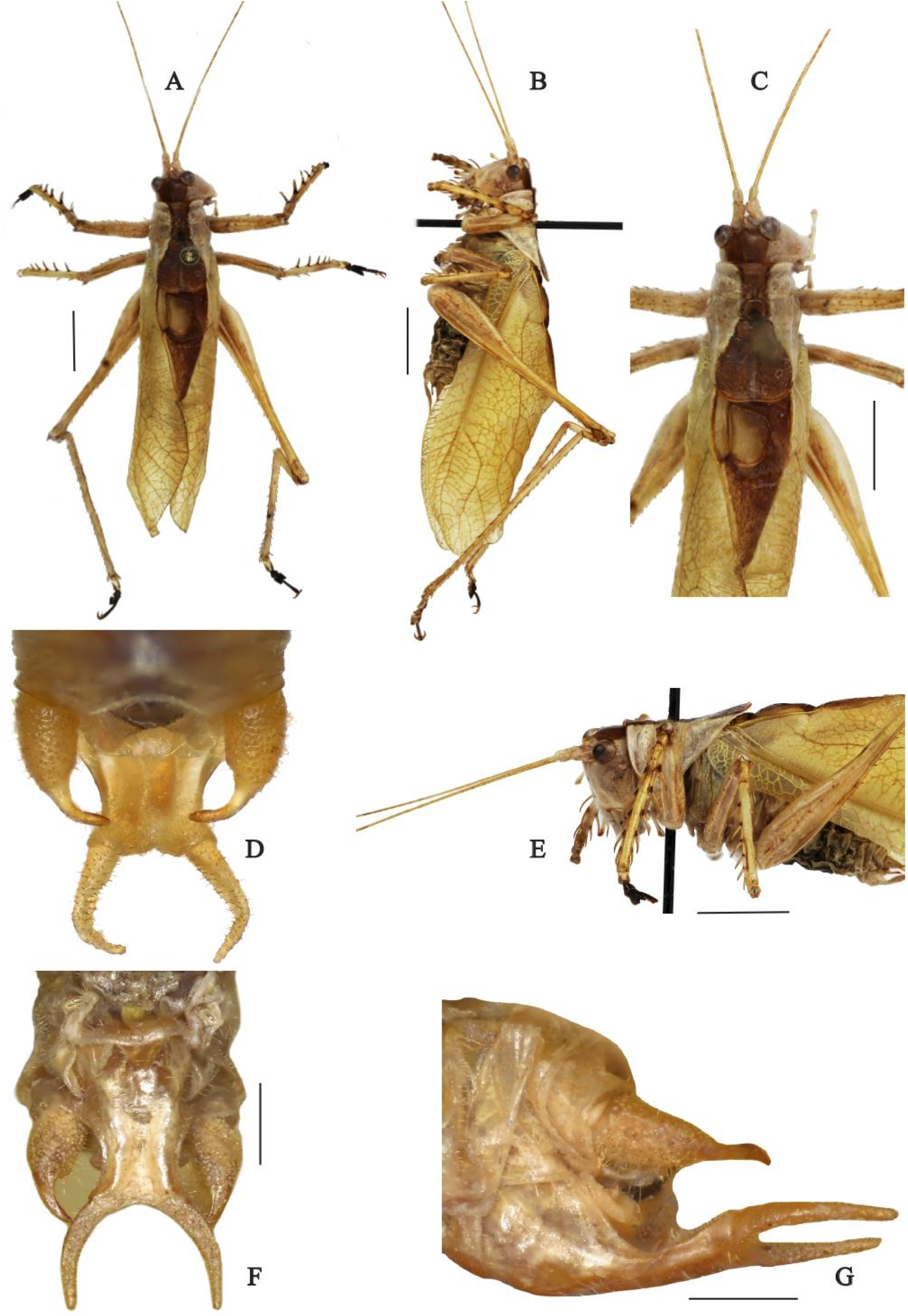
***Hexacentrus khasiensis*** male. (A-B) Dorsal and Lateral view of the specimen. Scale - 5mm, (C) Dorsal view of head and pronotum. Scale - 5mm, (D) Supra anal plate, (E) Front view of head and pronotum. Scale - 5mm, (F-G) Dorsal and Lateral view of sub genital plate, Scale - 1mm.

#### Male. Wings

Hindwings are shorter and hidden under tegmen; tegmen is medium size, broad in the middle, apical margin not oval in lateral view (Fig. 3B), reaching the middle of the hind tibia. Left tegmen has a long stridulatory file on the ventral side with 37-40 teeth; towards the origin of 1^st^ anal vein teeth are broader and situated at elevation, followed by slender teeth, middle part with broadest teeth, subsequent teeth gradually reducing in broadness and size. Mirror longer than wide, somewhat rectangular in shape. M vein slightly curved, CuP vein slightly bent in the middle, space between CuP and CuA slightly more than M and CuA, 1^st^ anal vein bearing stridulatory file inclined, length 1.3 mm, ventral side with 37-40 teeth, teeth density is 29.3; towards the origin of 1^st^ anal vein teeth are broader and situated at elevation, followed by slender teeth, middle part with broadest teeth, subsequent teeth gradually reducing in broadness and size (Fig. 14A).

#### Male. Genitalia

Supra anal plate is triangular in shape with a dorsal groove, slightly broad and short in length, the apex is round. Cerci robust basally long cylindrical in shape, then long tubercle on the internal side, then slowly narrowing to a long digitiform apical appendage with incurved apex (Fig. 3D). Sub genital plate elongate with a median longitudinal furrow, lateral ridges well developed, apical margin with deeply curved, styles are thick at base, apically thin and long, diverging basally and parallel apically (Fig. 3F).

#### Female

Small body size, cylindrical. **Pronotum**. Lateral lobe is somewhat straight in the posterior margin (Fig. 4F). **Wings**. FW narrow and short in length, reaching up to mid of ovipositor (Fig. 4B). **Female Genitalia**. Supra anal plate triangular with dorsal groove, short in length and broad basally, apex rounded (Fig. 4H). Cerci are conical, slightly upcurved, and apex diverging. The subgenital plate is triangular and wide (Fig. 4J). Apex with a median excision margin rounded apically. The ovipositor is broad at the base, straight, and mostly the same in width throughout the length except narrowing apically with a pointed end (Fig. 4I).

**Figure 4.**
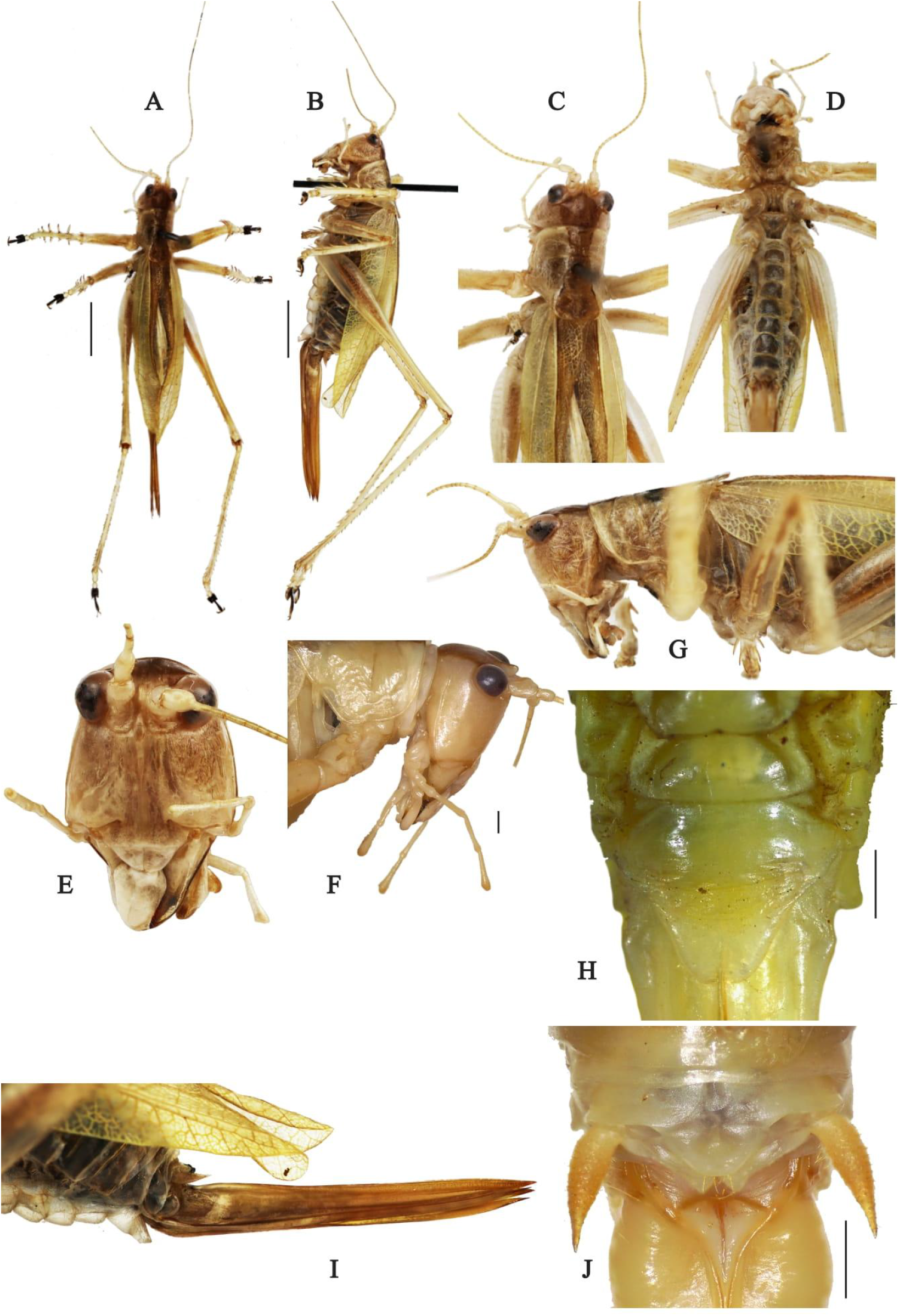
***Hexacentrus khasiensis*** female - (A-B) Dorsal and Lateral view of the specimen. Scale -5mm, (C) Dorsal view of head and pronotum, (D) Ventral view of Specimen, (E) Face, (F) Maxillary palpi. Scale - 1mm, (G) Lateral view of head and pronotum, (H) Dorsal view of sub genital plate. Scale - 1mm, (I) Ovipositor in lateral view, (J) Supra anal plate. Scale - 1mm.

**Figure 5.**
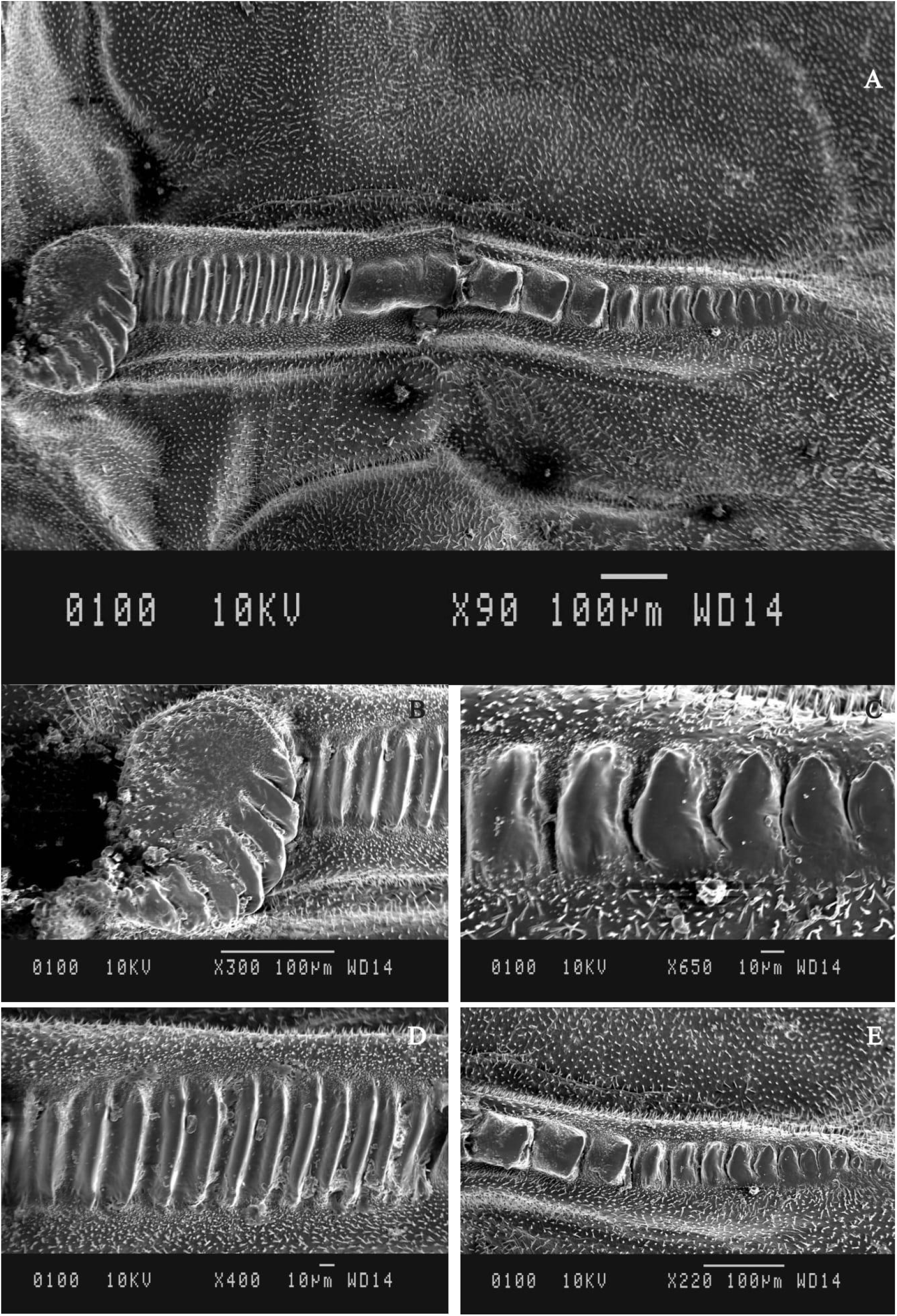
SEM images of the file structure of ***Hexacentrus khasiensis* sp. nov**. in different magnifications.

#### Coloration

Overall green when alive (Fig. 2B, 2D). Antennae with regularly spaced blackish brown annules. Occiput and vertex bearing thick rusty brown stripe covering most of the head in dorsal view. Pronotum DD bearing rusty brown stripe similar in width as of head up to half the length, much widened posteriorly. Male FW with stridulatory apparatus and apical field rusty brown, lateral part venation brown. The dorsal part of the female FW is rusty brown, lateral part venation brown. Inner side of FIII brown, knee joint between FIII and TIII brown, metatarsal and claw blackish brown.

### Acoustics

The *Hexacentrus khasiensis* **sp. nov**. call consists of a very interesting non-stop droning sound with two parts: the first part is a broadband continuous trill followed by the second part with an amplitude modulated trill (Fig. 6A). All measurements were made for at least 10 syllables per call portion per animal, for multiple animals. In the field, it sounds as if two different individuals are calling because of the continuous droning sound produced by opening and closing of the wing, but recordings in the lab verified that the call was produced by a singular individual.

**Figure 6.**
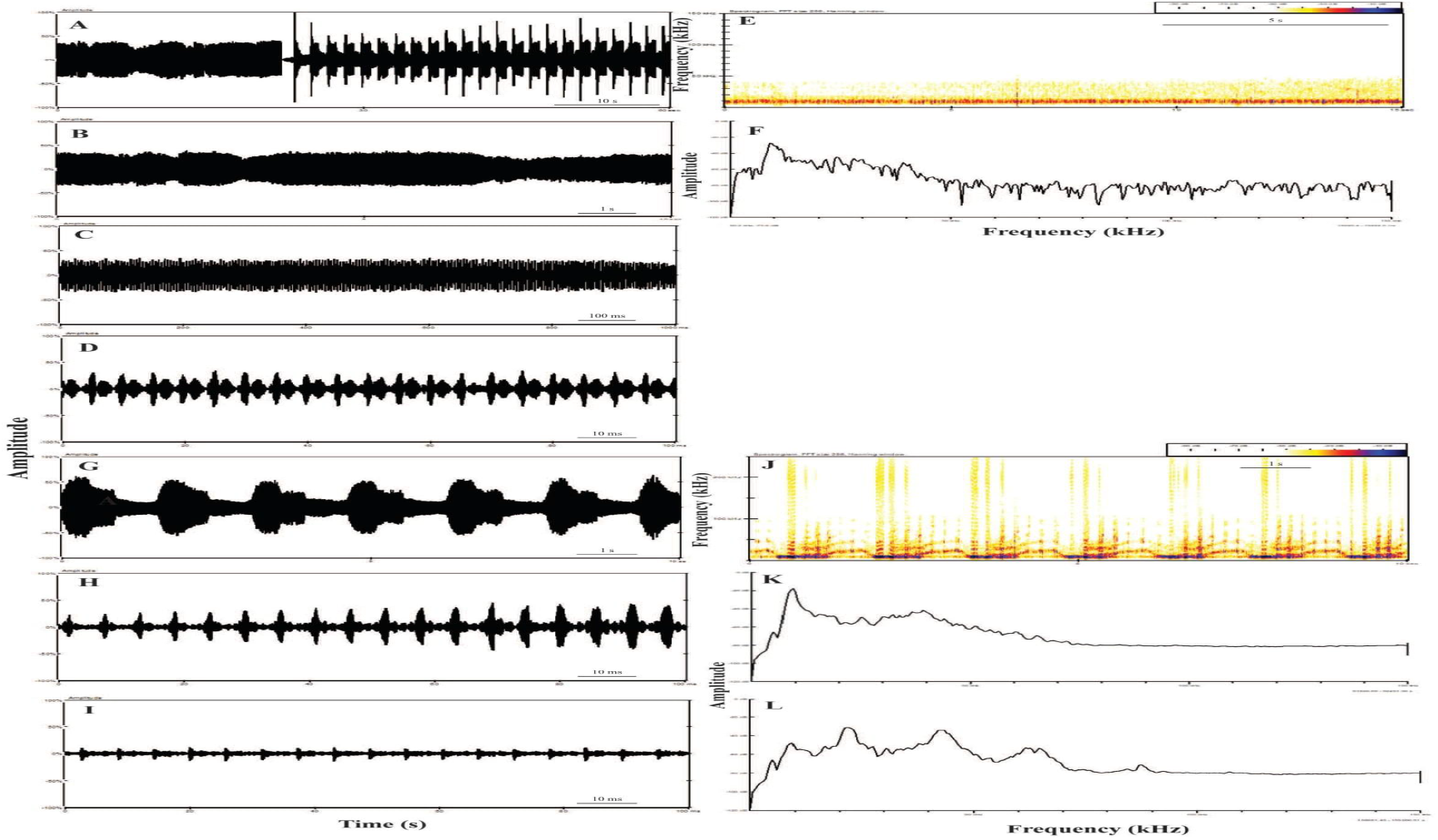
*Hexacentrus khasiensis* sp. nov. (A) Waveform of calling song, (B-D) Oscillogram of the first part of call at various time scales, (F) Spectrogram and (E) Power spectra of the first part of the call (G-I) Oscillogram of the second part of call at various time scales, (J) Spectrogram of the second part of call and Power spectra of the high amplitude syllables (K) and low amplitude syllables (L). ***Hexacentrus ashoka* Ghosh, Jaiswara & Rajaraman sp. nov**.

The call has these two distinct parts:

#### 1. First part

The first part of the call is a continuous buzz-like trill lasting 63.5-152.2 seconds (minimum and maximum duration measured in the field, **n=5**) (Fig. 6B), consisting of a series of evenly spaced syllables with a syllable repetition rate (SRR) of 172± 32.3. Each syllable is 2.2 ±0.5 ms in duration and the intersyllable interval is 3.9 ± 1 ms long (Fig. 6C-D). This part of the call has a peak frequency of 9 ± 0.1kHz and a bandwidth of 5.8 ± 1 kHz (Fig. 6E-F).

#### 2. Second part

After the first part, the call transitions to a series of amplitude and frequency-modulated groups of syllables, alternating between groups of high amplitude, high-frequency syllables, and portions with lower amplitude lower frequency syllables with slight frequency modulation over time. The syllable period and repetition rate at both sets of amplitudes remain the same at 4.7 ± 0.5ms and 214 syllables/ms respectively (n=10). The high amplitude syllable group lasts 741 ± 100 ms while the short amplitude group of syllables continues for a duration of 680 ± 90 ms (Fig. 6G-I). The high amplitude syllables have the same peak frequency of 9 ± 0.1kHz and bandwidth of 6.3 ± 1kHz, extending up to high frequencies beyond 250 kHz -we were limited by the spectral range of the recorder which only goes up to 250 kHz (Fig. 6J-K). The low amplitude syllables have a fundamental frequency of 9kHz, but there is a peak frequency band at 21±0.01 kHz with primary and secondary harmonics at 42kHz and 63kHz, respectively. Both the harmonics show some frequency modulation over time, but there is no discernible energy at frequencies above 65 kHz (Fig. 6L). The bandwidth of the low amplitude syllable part is 38 ± 2 kHz.

##### Type material

*Holotype*, **♂**, INDIA: Haryana, Sonipat, Rai, Rasoi ∼315 m a.s.l., VIII.2021,28° 56’ 48.78” N; 77° 6’ 5.19” E, AG_SO1_M3(ZSI). *Allotype*_**♀**, AG_SO1_F5, same locality as holotype (ZSI). *Paratypes* (2**♂**,3♀), same information as holotype, VII-IX.2020-2021: 1**♂**(AG_SO1_M1), 1**♂** ex-AG_SO1_M2 (BNHS); 2♀ (AG_SO1_F1, AG_SO1_F2), 1♀ ex-AG_SO1_F4 (BNHS).

##### Type locality

Ashoka University, Rai, Sonipat, Haryana, India.

##### Distribution

The species is currently known only from its type locality.

##### Etymology

The species is named after Ashoka University located in Sonipat, Haryana. The project was carried out with the support and research grant funding provided by Ashoka University. The University is in turn named after the pluralistic and peace-loving Ashoka, after whom this species has been named.

##### Diagnosis

The species can be diagnosed from *H. khaisiensis* **sp. nov**. with pronotum anterior margin slightly concave, pronotal disc flat, posteriorly expanded, broadly rounded, posterior margin concave with slight median excision. Longer wings cover the whole abdomen and M vein is much more curved, thick distally, CuP vein somewhat straight, space between CuP and CuA slightly same as between M and CuA, 1^st^ anal vein bearing stridulatory file less inclined, ventral side with 40 teeth (Fig.14B,9). Cerci are cylindrical in shape at the base; it is broad with a tubercle on the internal side then narrowing to small digitiform apical appendages with strongly incurved apex.

##### Description

Mostly similar to *H. khaisiensis* **sp. nov**. Except for the following characters.

**Male**. Body slender in shape, medium-sized (Fig. 7A-B). **Head**. The fastigium of vertex is triangular, narrow, laterally compressed with a median longitudinal sulcus, and apex round in lateral view. Base of the fastigium vertex is almost the same width as the scapus. Fastigium of vertex is separated by the fastigium of fons by a farrow. Eyes are globular (Fig. 7C-E).

**Figure 7.**
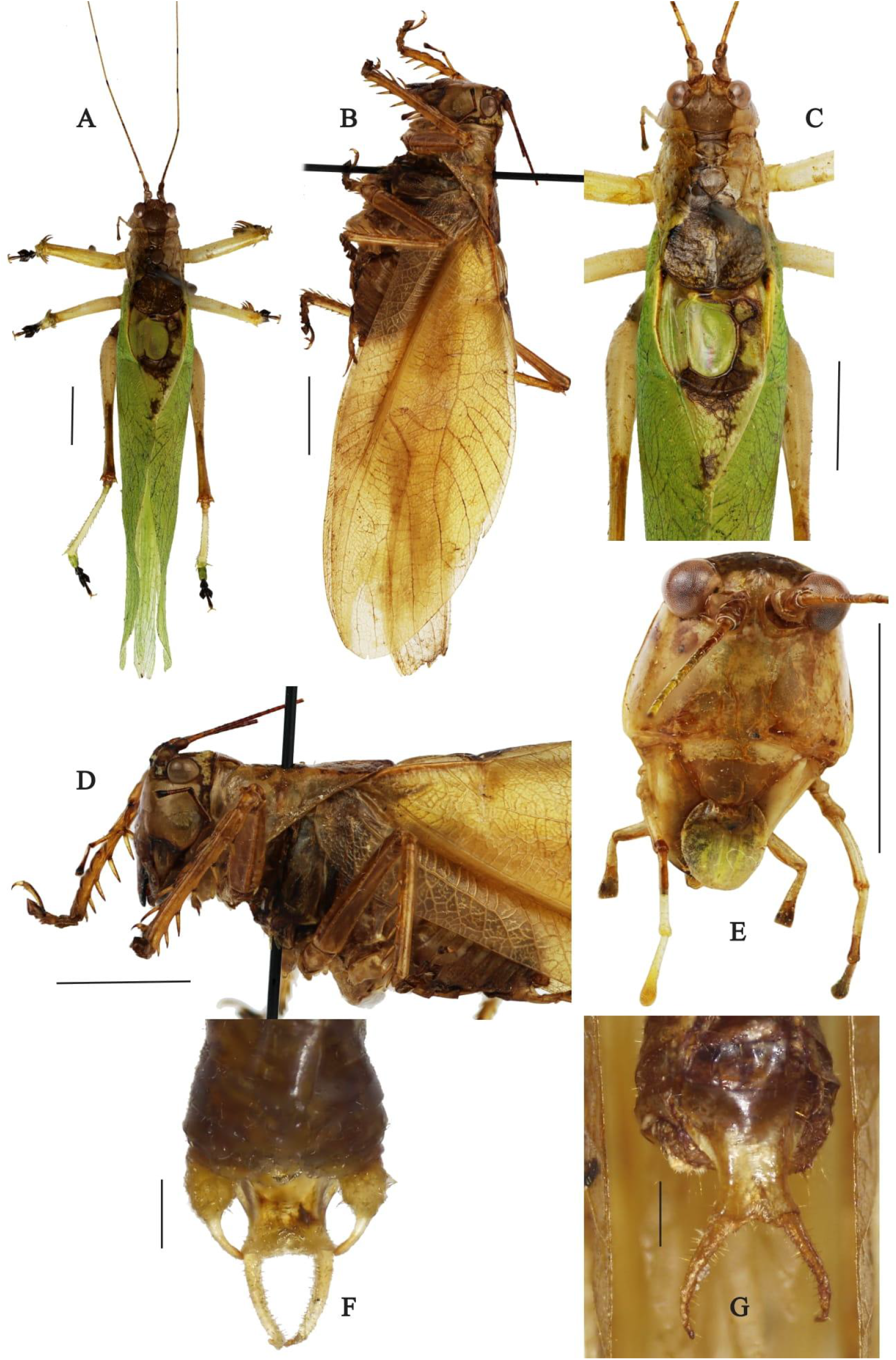
***Hexacentrus ashoka*** Male- (A-B) Dorsal and Lateral view of the specimen. Scale - 5mm, (C-D) Dorsal and Lateral view of head and pronotum. Scale - 5mm, (E) Face. Scale - 5mm, (F) Supra anal plate. Scale - 1mm, (G) Sub genital plate dorsal view. Scale - 1mm.

##### Pronotum

Saddle shaped, longer than wide and expanded in posterior region; pronotum has three transverse sulci, slightly depressed behind the first transverse sulcus, the third transverse sulcus is restricted to the disc, “U” shaped sulcus presents between second and third transverse sulci, anterior margin slightly concave with no excision present. An hourglass-like brown band is present in the pronotum. Median carina is faintly visible (Fig. 7C). Pronotal pronotal disc is flat, posteriorly expanded, broadly rounded, posterior margin concave with slight median excision. Lateral margin of pronotum descending, longer than high, with the humeral sinus absent, anterior angle is round and anterior margin shorter in length than posterior margin from lateral view, posterior angle is obtuse. Thoracic auditory spiracle large, oval, slightly hidden under lateral lobe of pronotum (Fig. 7D).

##### Wings

Tegmina is slightly shorter or the same as the length of hind wings. Tegmina is long and crosses the mid tibiae of the hind leg. Male tegmina is long and broad in the middle, the apex is narrow and round in lateral view (Fig. 7B). The Left-wing has the stridulatory file on the ventral side, shorter in length. It is broader at the tip and narrows towards the center. Tooth length is different in different parts of the file, widest in the beginning. The stridulatory file has 40 teeth (Fig. 9).

##### Legs

Fore coxae have a forward outward projecting spine. FI with 4 sub apical spurs and 1 apical spur, 16 spines in between sub-apical spurs on the inner margin, and approximately 27 spines with no sub apical and apical spur on the outer margin.

TI dorsally unarmed and ventrally armed with 6 long sub apical spurs inwardly bent, 1^st^ sub apical spur slightly shorter than 2^nd^ sub apical spur and then decreasing in length apically on both margins. TII is dorsally armed with two sub apical spurs, sometimes absent, ventrally armed with 6 sub apical spurs on inner and outer margins. TIII dorsally armed with 24-36 spines on the inner margin and 27-35 spines on the outer margin, TIII ventrally armed with 10-15 sub apical spurs increasing in length and density each on the outer margin and 8-15 sub apical spurs increasing in length and density towards tarsus.

##### Male Genitalia

Supra anal plate triangular in shape with dorsal groove. Cerci are cylindrical in shape at the base; it is broad with a tubercle internal side then narrowing to small digitiform apical appendages with strongly incurved apex (Fig.7F). Sub genital plate is long with elongated, wide longitudinal median furrow, and lateral ridges are well developed, apical margin with wide V-shaped excision, styles curved, slightly thick at the base than apex, diverging basally and slightly converging apically (Fig. 7G).

### Female

#### Pronotum

From the dorsal side posterior lobe of the female pronotum is less wide and half-rounded than the males (Fig. 8C). **Wings**. Hind wings and tegmina are the same in length. Tegmina is long and narrow same width throughout, the apex is rounded (Fig. 8B).

**Figure 8.**
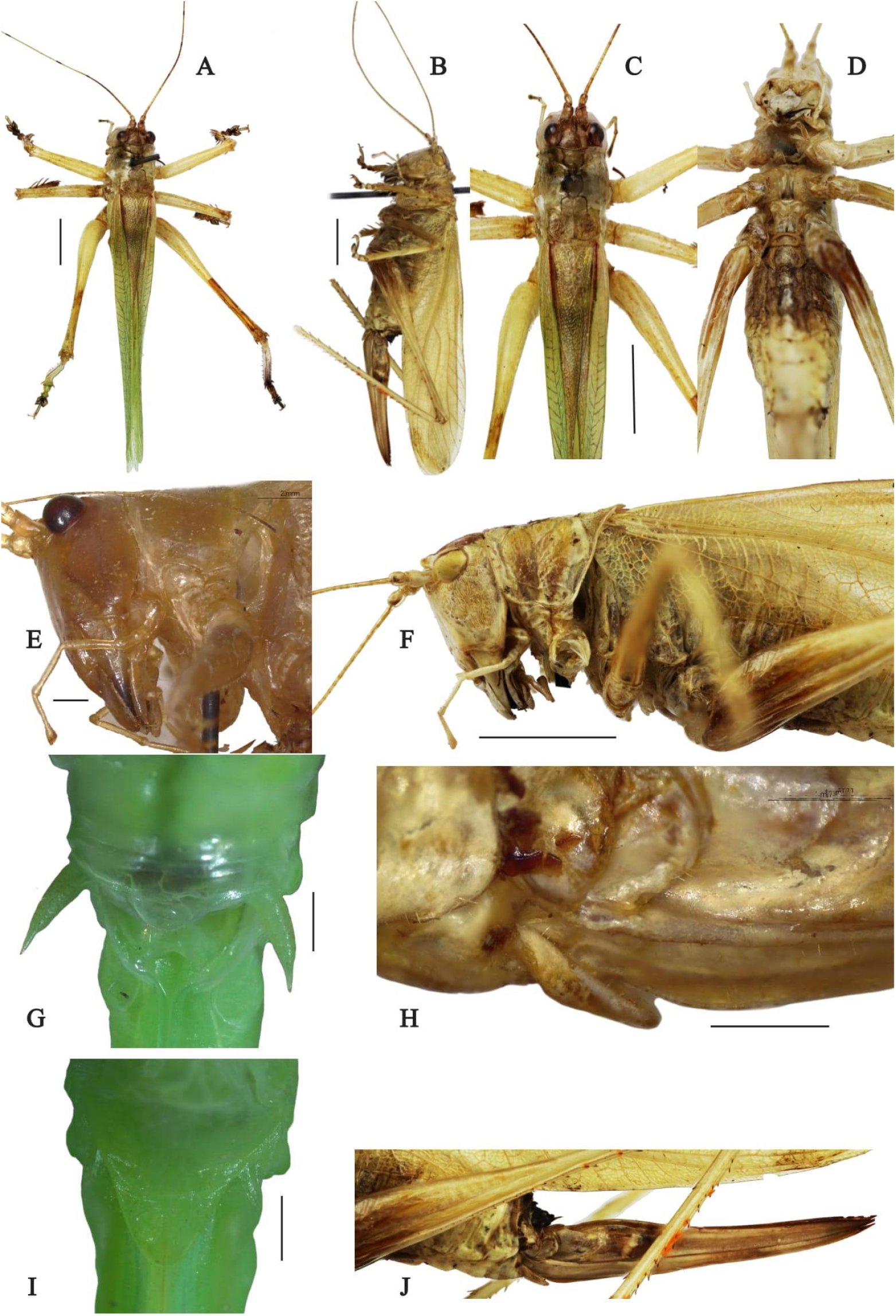
***Hexacentrus ashoka*** Female -(A-B) Dorsal and Lateral view of the specimen. Scale - 5mm, (C) Dorsal view of head and pronotum. Scale -5mm, (D) Ventral view of Specimen, (E) Maxillary palpi. Scale - 1mm, (F) Lateral view of head and pronotum. Scale - 5mm, (G) Supra anal plate. Scale - 1mm, (H-I) Lateral and Dorsal view of sub genital plate. Scale - 1mm, (J) Ovipositor in lateral view.

**Figure 9.**
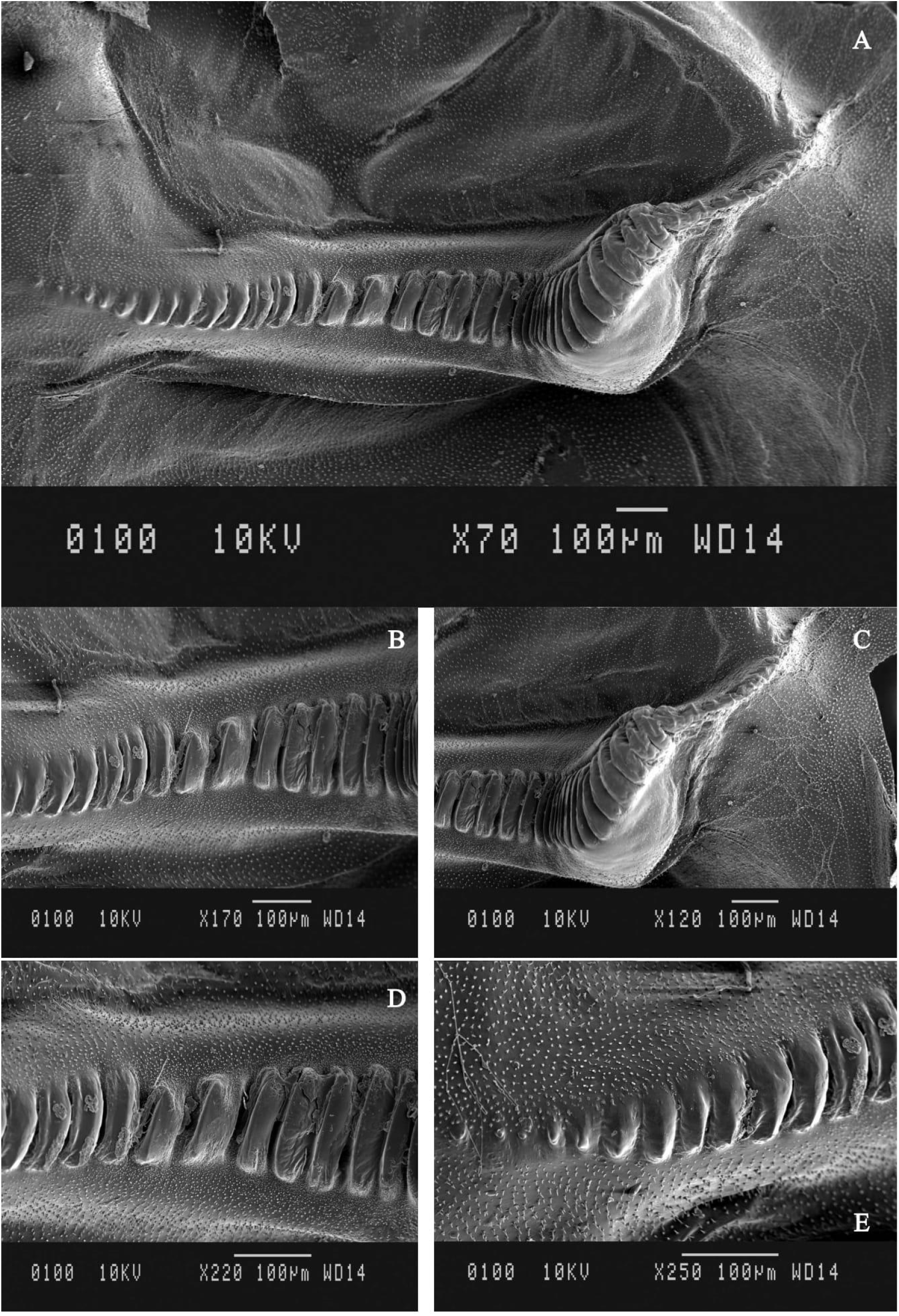
SEM images of the file structure of ***Hexacentrus ashoka* sp. nov** in different magnifications.

#### Female Genitalia

Supra anal plate triangular with dorsal groove, short in length and broad basally, apex round. Sub genital plate is triangular, and small, with a dorsal groove, length and width are the same, apex is bifurcated and rounded apically (Fig. 8I). Cerci are robust, upcurved, straight, thick at the base narrow at the apex, with smooth conical slightly inward appendages (Fig. 8G). Ovipositor dagger shaped, the base is medium-sized, near the middle is widest, gradually narrowing towards the apex, apex pointed, serrated apically (Fig. 8J)

#### Coloration

Overall green when alive (Fig.2A). Antennae with regularly spaced blackish brown annules. Occiput and vertex bearing thick rusty brown stripe covering most of the head in dorsal view. Pronotum DD bearing rusty brown stripe less in width as of head up to half the length much widened posteriorly. Male FW area around the mirror and apical field rusty brown, lateral part venation brown. The dorsal part of the female FW is rusty brown, lateral part venation brown. FIII brown half the length, knee joint between FIII and TIII brown, metatarsal and claw blackish.

### Acoustic measurements

The *Hexacentrus ashoka* call consists of two sections (Fig. 10A). The first part of the call consists of a group of low-amplitude chirps with variable chirp duration: the earlier chirps are longer (337 ± 1.7ms for the first five chirps from this portion averaged across 3 individuals) and the later chirps are quicker (261 ±18 ms for the last five chirps from this portion averaged across 3 individuals). There is also considerable variability in the inter-chirp interval, which also reduces over time (317 ± 10ms for the first five chirps, 187.2 ± 13ms for the last five, n = 3 individuals). The second part has a group of higher amplitude and well-defined sharp chirps that sound like short buzzing sounds (Fig. 10B-D). Each chirp ends with a quick single syllable tick of 2 ± 0 ms length. The high amplitude chirps have groups of 80.8 ± 2.13 syllables for a total duration of 194 ± 50 ms (n=7). Each syllable is 1.7 ± 1 ms in length with inter syllable duration of 1.2ms. The SRR is 340 ± 18. The spectrogram is broadband with a bandwidth of 5 ± 0.8kHz and a peak frequency of 8.9± 0.13kHz (Fig. 10E-F).

**Figure 10.**
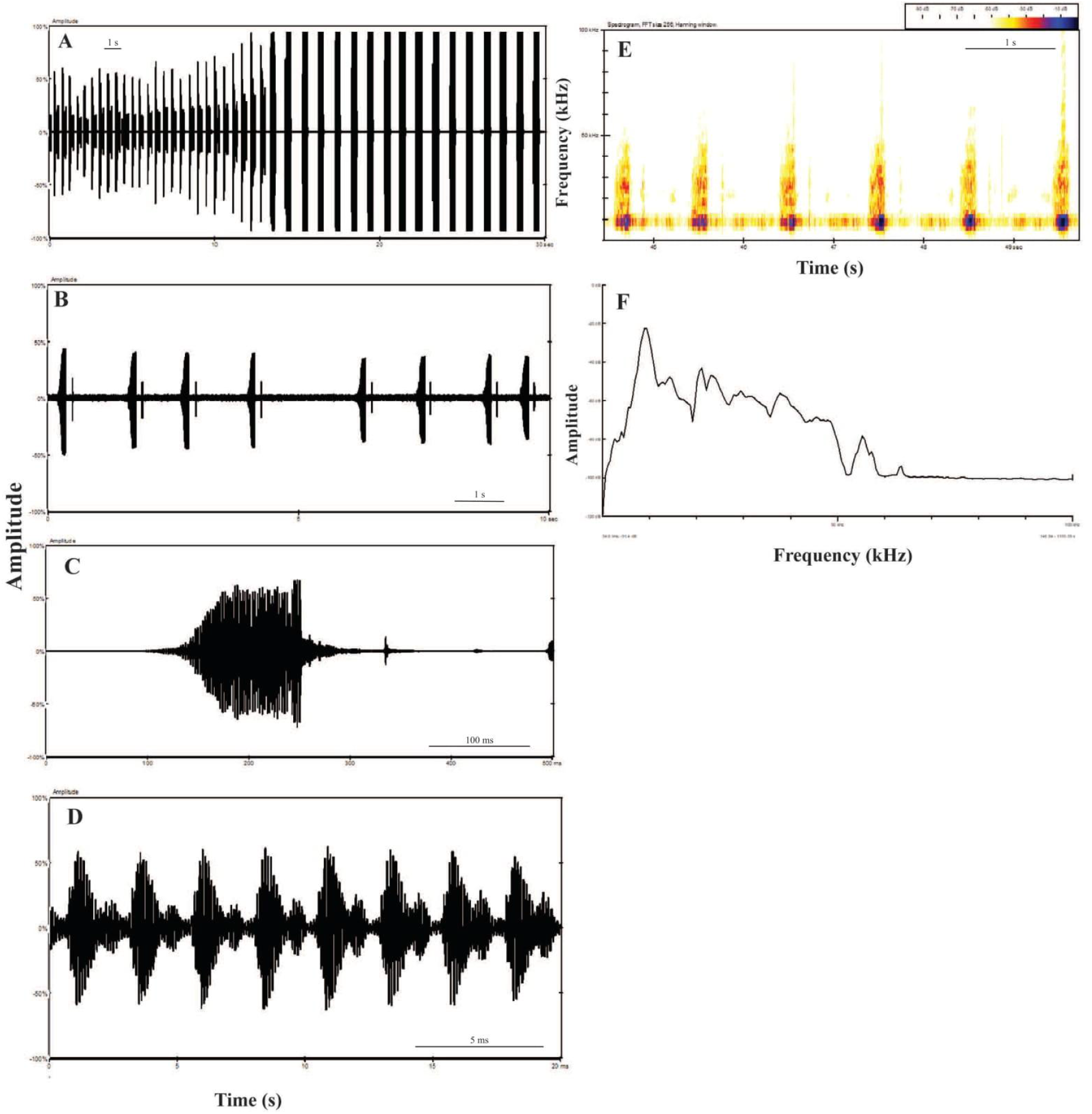
***Hexacentrus ashoka* sp. nov**. (A) Waveform of whole calling song, (B-D) Oscillogram of the call at various time scales, (E) Spectrogram, and (F) Power spectra of the calling song.

### *Hexacentrus tiddae* Ghosh, Jaiswara, Monaal & Rajaraman sp. nov

#### Type Material

Holotype, **♂, INDIA:** Haryana, Sonipat, Rai, Aswarpur, ∼315 m a.s.l., 10.VII.2022, 28° 56’ 48.78’’ N; 77° 6’ 5.19” E, AG_SO2_M1(ZSI). Paratype-1**♂, the** same information as holotype, VII-2022 ex-AG_SO2_M2, without left forewing (BNHS).

#### Type locality

Ashoka University, Aswarpur, Sonipat, Haryana, India.

#### Distribution

The species is known from its type locality only for now.

#### Etymology

‘tiddae’ in Hindi and Haryanvi refers to Orthoptera - it is used for locusts and crickets.

#### Diagnosis and Description

The species is somewhat similar to *H. ashoka, H. bifurcata* & *H. japonicus* and differs in the following characters. Body medium size and slender (Fig. 11A-B). **Head**. Face longer than wider; almost double. Antennae without any short dark bands both apically and throughout the length. Fastigium of the vertex laterally much more compressed, apically tapering and triangular in shape in dorsal view with pointy appearance apically. Scapes cylindrical with slight bulge at apex in dorsal view, scapes almost 3 times as wide as fastigium (Fig. 11C & 11F). Cheeks are slightly broad.

**Figure 11.**
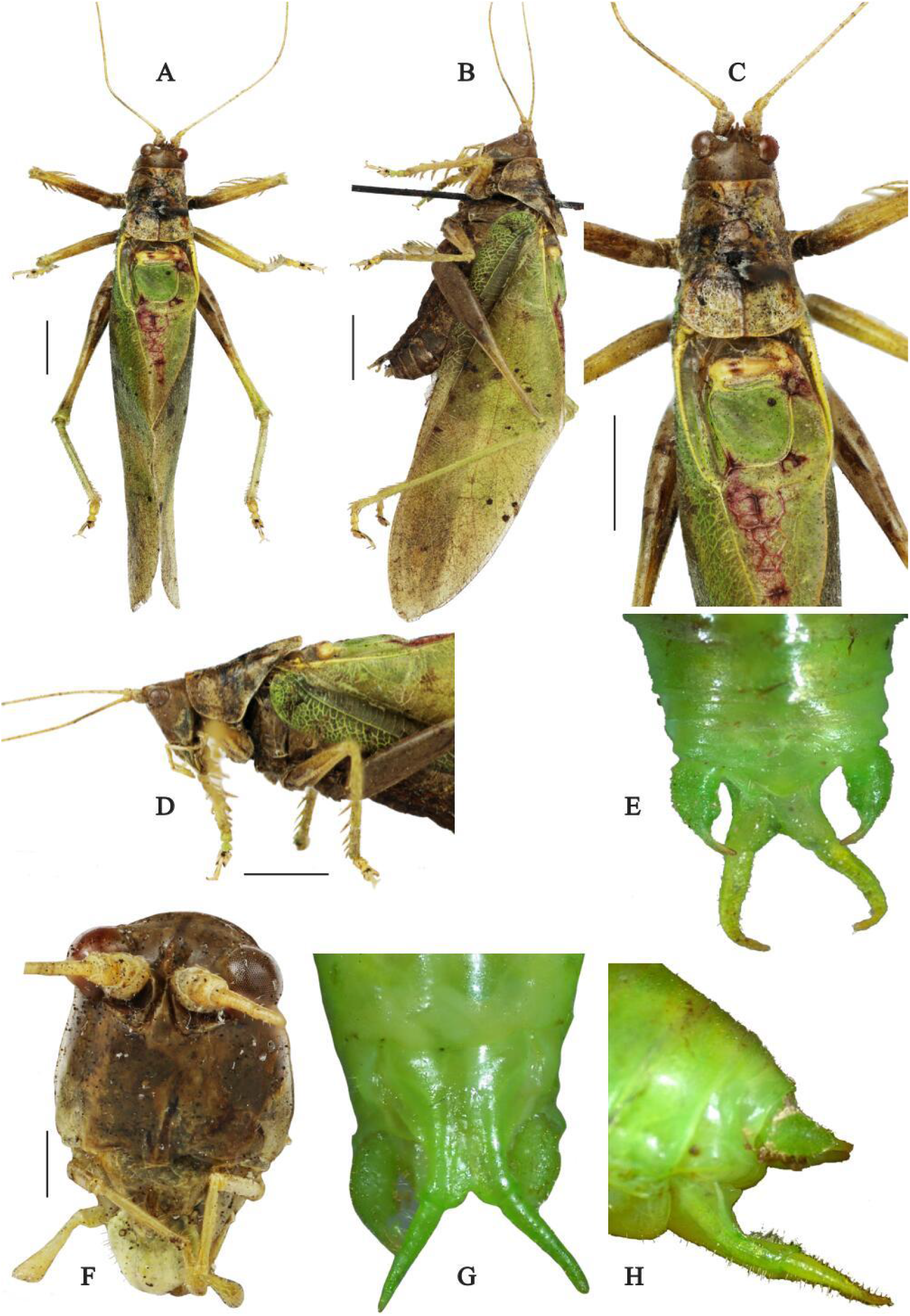
***Hexacentrus tiddae*** male - (A-B) Dorsal and Lateral view of the specimen. Scale - 5mm, (C) Dorsal view of head and pronotum. Scale -5mm, (D) Lateral view of head and pronotum. Scale - 5mm (E) Supra anal plate. Scale - 1mm, (F) Face. Scale - 5mm, (G-H) Lateral and Dorsal view of subgenital plate. Scale - 1mm.

**Figure 12.**
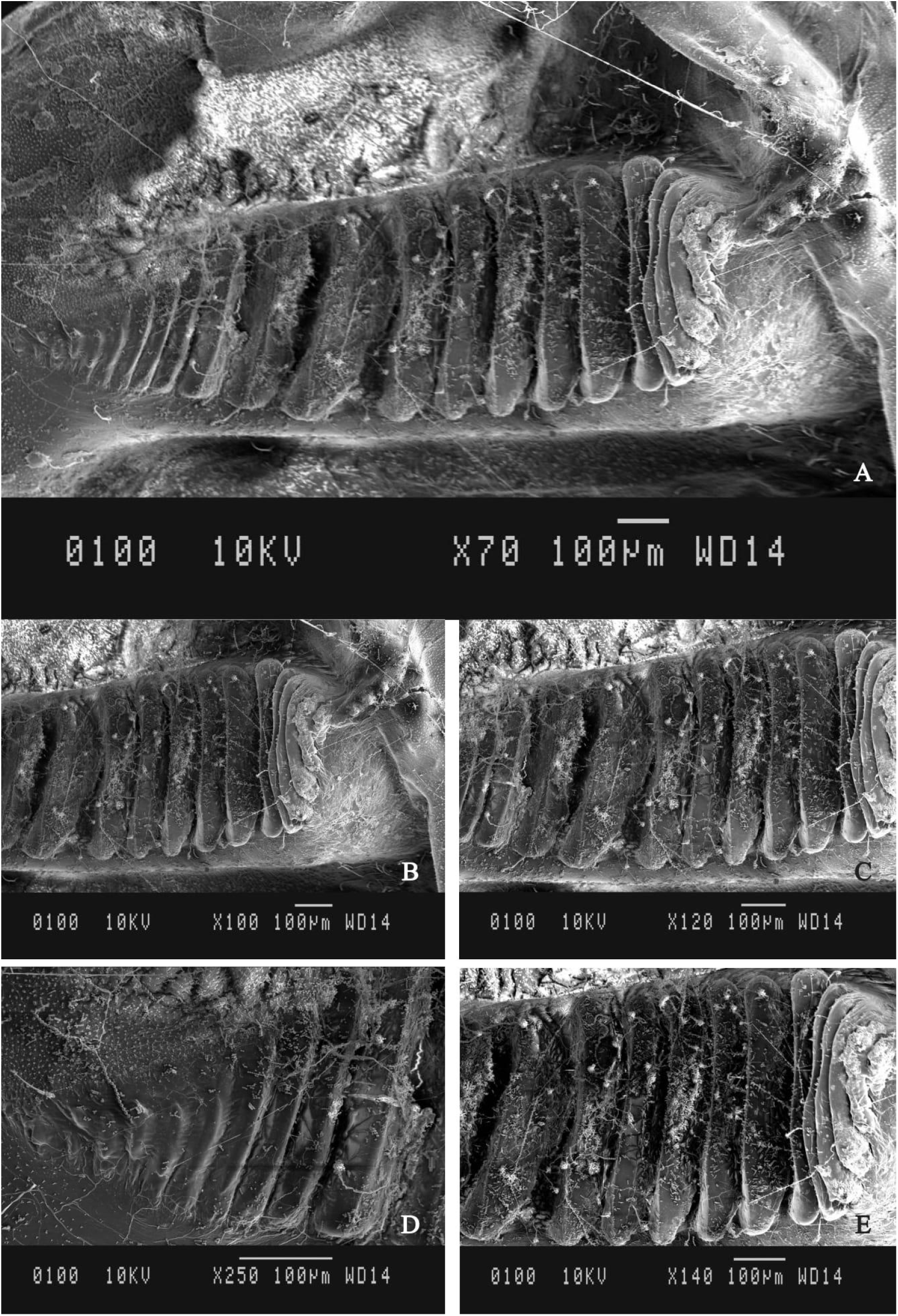
SEM images of the file structure of ***Hexacentrus tiddae* sp. nov**. in different magnifications.

#### Pronotum

Pronotum is longer than wide, flat with a slightly upward curve posteriorly, longer than broad dorsally and extended posteriorly. The anterior and posterior margins slightly concave; the hourglass-like band covers the whole pronotum (Fig. 11C-D).

#### Legs

FI dorsally unarmed and ventrally armed with 18-30 spines on inner and outer margins, 1 apical spine on lateral margin; TI has a tympanum on both the inner and outer side, appearing ellipsoidal in dorsal view, inner side tympanum slightly longer than outer tympanum; TI dorsally unarmed and ventrally armed with 6 subapical spurs on each side, both inner and outer margin with no spines; FII dorsally unarmed and ventrally armed with 18-22 spines on inner and outer margins, 2 apical spines in the lateral margin; TII dorsally unarmed and ventrally armed with 6 sub apical spurs on each side, both inner and outer margin with no spines; FIII dorsally unarmed and ventrally armed with 39-52 spines both or inner and outer margin; TIII dorsally armed with 34-36 spines on both inner and outer margins and ventrally armed with 11-14 sub apical spurs on both inner and outer margins, one subapical spur are present in the center.

#### Male. Wings

Hindwings are almost as long as tegmina, hidden under tegmen; tegmina is long and crosses the tibiae of the hind leg. Male tegmina are long, broad in the middle, and somewhat triangular at the apex. M vein straight, thick distally, CuP vein bent distally, space between CuP and CuA less than between M and CuA, 1^st^ anal vein bearing stridulatory file somewhat straight, bent in the middle, ventral side with 20-23 teeth (Fig.14C,12).

#### Male Genitalia

Supra anal plate triangular in shape with dorsal groove. Slightly broad and short in length the apex is slightly pointed. Cerci are cylindrical in shape at the base; it is broad with a tubercle on the internal side, then abruptly narrows to small digitiform apical appendages with strongly incurved apex (Fig.11E). Sub-genital plate elongates with a median longitudinal furrow, lateral ridges well developed, apical margin with prominent ‘V’-shaped bifurcation (Fig.11G). Styles are slightly thicker at the base than apically, both the styles are diverging from each other, and the apex is slightly rounded and inward curved (Fig. 11G-H).

### Acoustic data

*Hexacentrus* ***tiddae*** produces a regular amplitude modulated pattern of continuous syllables, alternating between longer, higher amplitude syllables and quicker, low amplitude syllables. All these syllables have the same broadband spectral structure. The peak frequency is 9.4±3.6 kHz (Fig. 13D-E) and the bandwidth is 40 ± 3.2 kHz (Fig. 13D-E). The low amplitude syllables last for 0.9 ± 0.4 ms, with a total syllable period of 1.61 ±0.6 ms (Fig. 13B-13C), and SRR of 715 ± 156 (n=3 animals, 10 syllables of each kind per animal). The high amplitude syllables occur regularly, separated by an average interval of 846.25 ± 46 ms. The high amplitude syllable duration is 2.33±0.7 ms with a syllable period of 1.71 ±0.4ms and SRR of 463 ± 120.2 (Fig. 13C) (n=3 animals, 10 syllables per animal). The juxtaposition of the high amplitude buzzing bursts and the low amplitude background trill sounds almost like two simultaneous callers.

**Figure 13.**
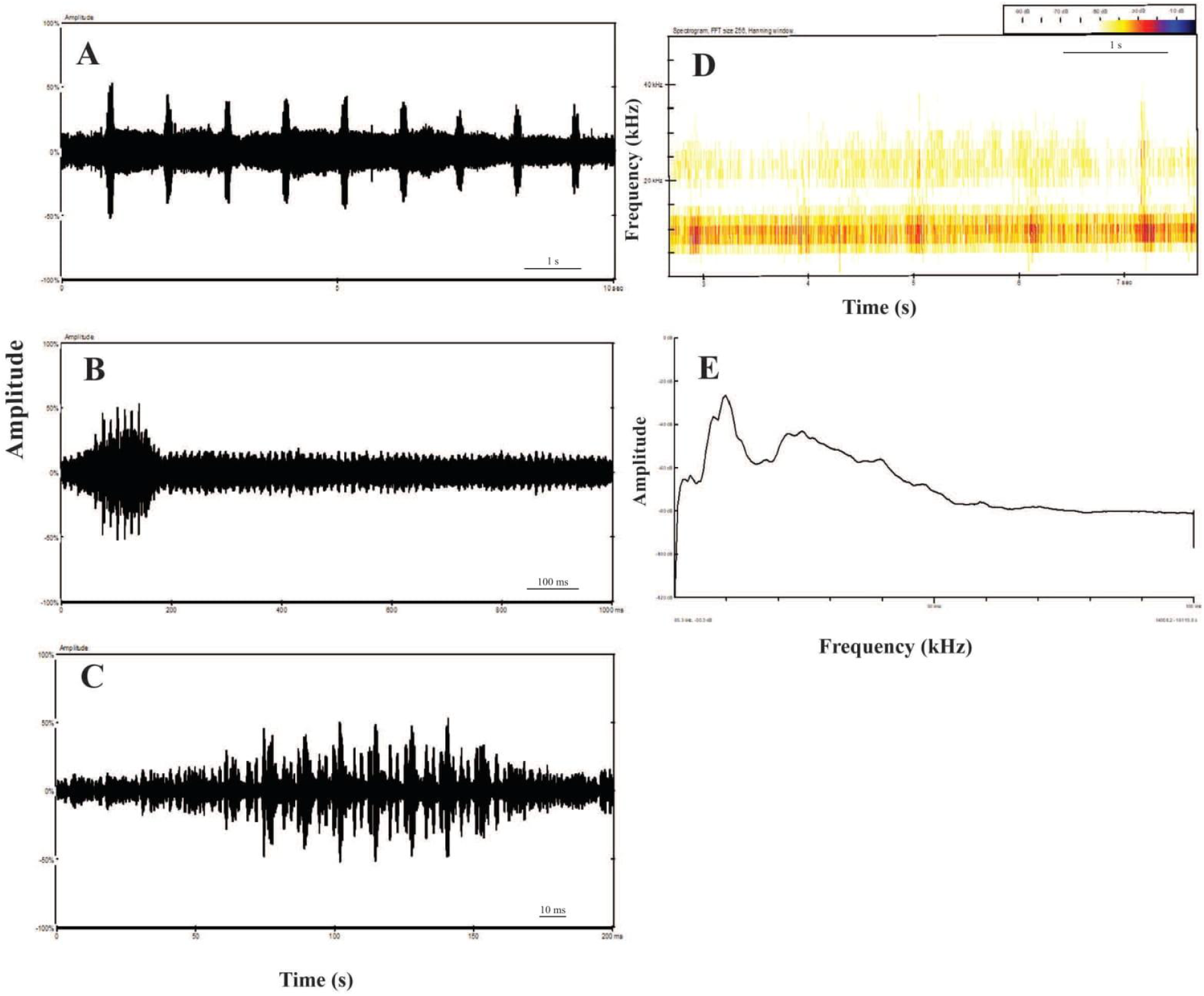
*Hexacentrus tiddae* sp. nov. (A-C) Oscillogram of the call at various time scales, (D) Spectrogram, and (E) Power spectra of the calling song.

**Figure 14.**
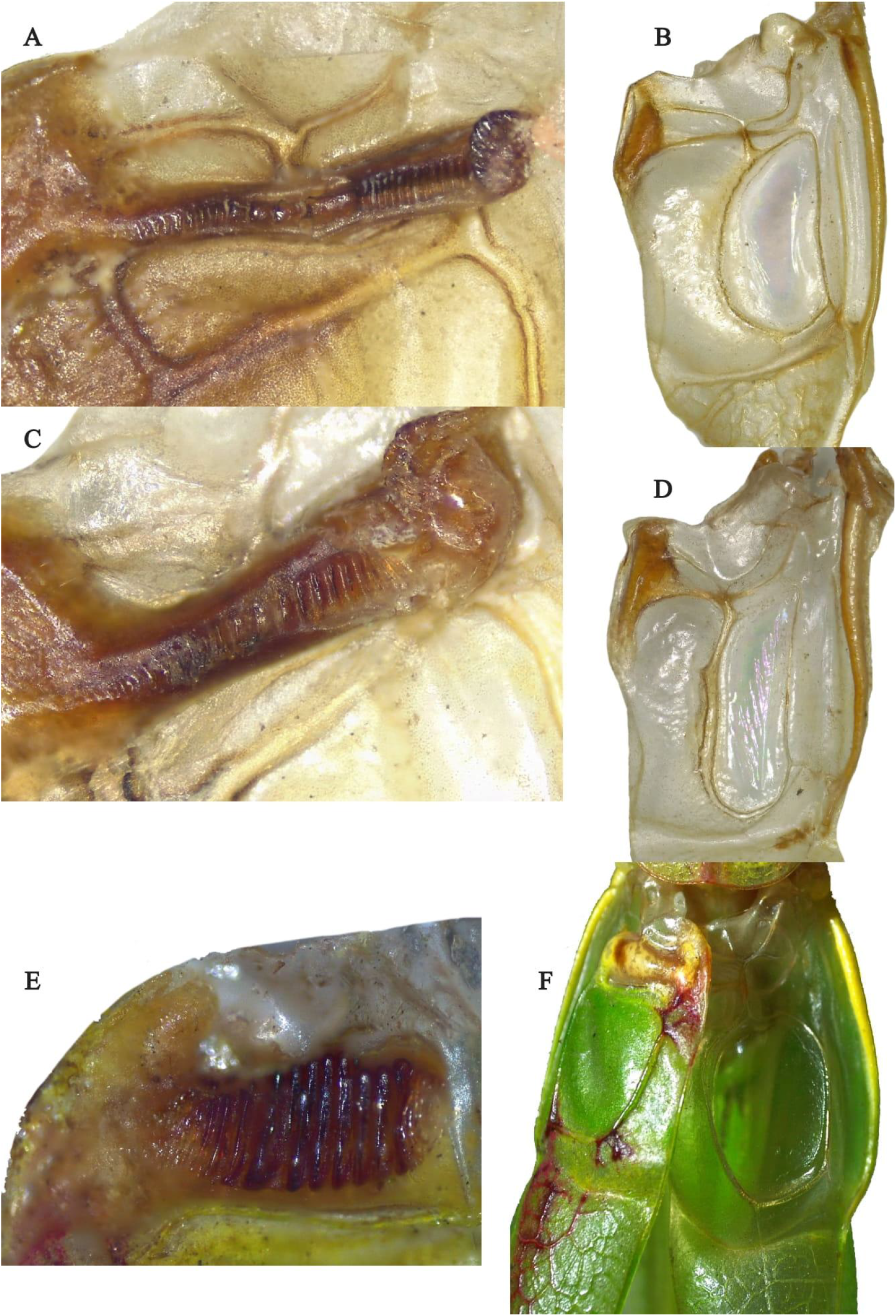
Stridulatory apparatus structure - ventral side of left wing with file and scraper on dorsal side of right wing (A-B) of ***H*.*khasiensis*** ; (C-D) of ***H. ashoka***; (E-F) of ***H*.*tiddae***.

## Acknowledgments

We acknowledge the DST-CRG scheme and the Ashoka University Annual Research Grant for funding AG and BKR, and the DST-INSPIRE Faculty scheme Government of India for funding RJ to conduct this study. Thanks, are also due to the Chairman, Department of Zoology, Panjab University, Chandigarh, India, for providing infrastructural facilities, to Dr. Sudipta Tung’s lab for sharing lab space and microscopy equipment, Dr. Imroze Khan’s lab group for supplying larvae to feed these predatory katydids, and to Dr. Shivani Krishna and Dr. Manjari Jain for academic guidance. Ashique Rahman, Deb Konar, Riban war, Wanshan, Shweta Prasad, Vikrant Menia, Prabhat Sharma, Pranya Prakash, Vaishnavi Agarwal, and Damayanti Dasgupta all contributed to the fieldwork. We acknowledge the immense support from the Meghalaya Forest Department and the Khasi community people from Meghalaya regarding permission to work in the Khasi hills.

